# MetaPrism: A Toolkit for Joint Taxa/Gene Analysis of Metagenomic Sequencing Data

**DOI:** 10.1101/664748

**Authors:** Jiwoong Kim, Shuang Jiang, Guanghua Xiao, Yang Xie, Dajiang J. Liu, Qiwei Li, Andrew Koh, Xiaowei Zhan

## Abstract

**Background:** In microbiome research, metagenomic sequencing generates enormous amounts of data. These data are typically classified into taxa for taxonomy analysis, or into genes for functional analysis. However, a joint analysis where the reads are classified into taxa-specific genes is often overlooked.

**Result:** To enable the analysis of this biologically meaningful feature, we developed a novel bioinformatic toolkit, MetaPrism, which can analyze sequence reads for a set of joint taxa/gene analyses: 1) classify sequence reads and estimate the abundances for taxa-specific genes; 2) tabularize and visualize taxa-specific gene abundances; 3) compare the abundances between groups, and 4) build prediction models for clinical outcome. We illustrated these functions using a published microbiome metagenomics dataset from patients treated with immune checkpoint inhibitor therapy and showed the joint features can serve as potential biomarkers to predict therapeutic responses.

**Conclusions:** MetaPrism is a toolkit for joint taxa and gene analysis. It offers biological insights on the taxa-specific genes on top of the taxa-alone or gene-alone analysis.

MetaPrism is open source software and freely available at https://github.com/jiwoongbio/MetaPrism. The example script to reproduce the manuscript is also provided in the above code repository.

## Introduction

The human microbiome consists of ∼39 trillion bacteria and influences host health. Recently, the use of metagenomic sequencing has become increasingly popular as a more unbiased approach to gut microbiome profiling as compared to 16S rRNA sequencing. A common approach to comparing differences in the gut microbiome between groups (cases and controls) is to identify significant differences in either taxa or microbial genes. Several popular bioinformatic tools have been developed for this purpose, including MetaPhlAn2 [1], Kraken [2], HUMAnN2 [3], and FMAP [4] (**Table S1**). However, these tools analyze either taxonomic abundances (taxonomic profiling) or gene abundances (function profiling) separately. As each microorganism carries its own genes, taxonomic and functional profiling results are not intrinsically independent. In fact, recent discoveries demonstrated that taxon-specific genes have a causative role in disease progression and treatment responses. For example, Duan et al. found that a specific *Enterococcus faeclis* carrying the cytolysin gene promotes alcoholic liver disease[5]. Simms-Waldrip et al. found that the antibiotic resistance genes in the graft-versus-host-disease patients are enriched for *Klebsiella*[6]. Therefore, joint analysis, where taxonomy and functional features are analyzed together, could provide useful biological and clinical insights [7]. However, bioinformatics tools for joint analyses are comparatively lacking.

Our innovation in this manuscript is to define and utilize joint taxa/gene features via bioinformatics approach, with the goal of offering biologically interpretable findings. For example, our method characterizes the genes discovered for each species. This allows to quantitative analysis of this species-specific gene, which is usually not readily available. Our approach is initiated from *de novo* assembled contigs which are both taxonomically and functionally annotated. Our simulations showed this method can accurately detect bacterial species and their carried genes. In a recent review article[7], Langille prompted that understanding the gene contents at species level can offer better interpretation than using the taxon or gene content alone, and potentially provide better prediction outcomes. This confirmed that the joint feature is useful for general microbiome studies. Our tool provided these joint features as the first step for a wide range of downstream analysis tasks. For example, we demonstrated that the quantity of taxa-specific gene abundances is a potentially useful biomarker to predict the immunotherapy responses.

To facilitate joint analysis, we developed MetaPrism, a novel bioinformatics tool to (1) classify metagenomic sequence reads into both taxa and gene level, (2) normalize the taxa-specific gene abundances within samples, (3) tabularize or visualize these joint features, (4) perform comparative microbiome studies, and (5) build prediction models for clinical outcomes. MetaPrism is open-sourced and is available at https://github.com/jiwoongbio/MetaPrism. Given the advantages of joint analysis, MetaPrism is a useful tool for microbiome metagenomic sequence studies.

## Material and Methods

MetaPrism is a toolkit for joint analysis tasks. At its core, MetaPrism will infer the taxa and gene for each metagenome sequence read. One approach is to align each read to bacterial nucleotide reference genomes to obtain its taxonomy and align it to a protein database to obtain its gene functions. However, this approach is technical challenging: due to the short lengths of Illumina sequence reads and the high sequence similarities between bacteria genomes, alignment of short reads is not feasible. We thus developed a novel algorithm (**Figure 1A**) in an integrated toolkit (**Figure 1B**) to tackle this challenge.

**Figure 1.**
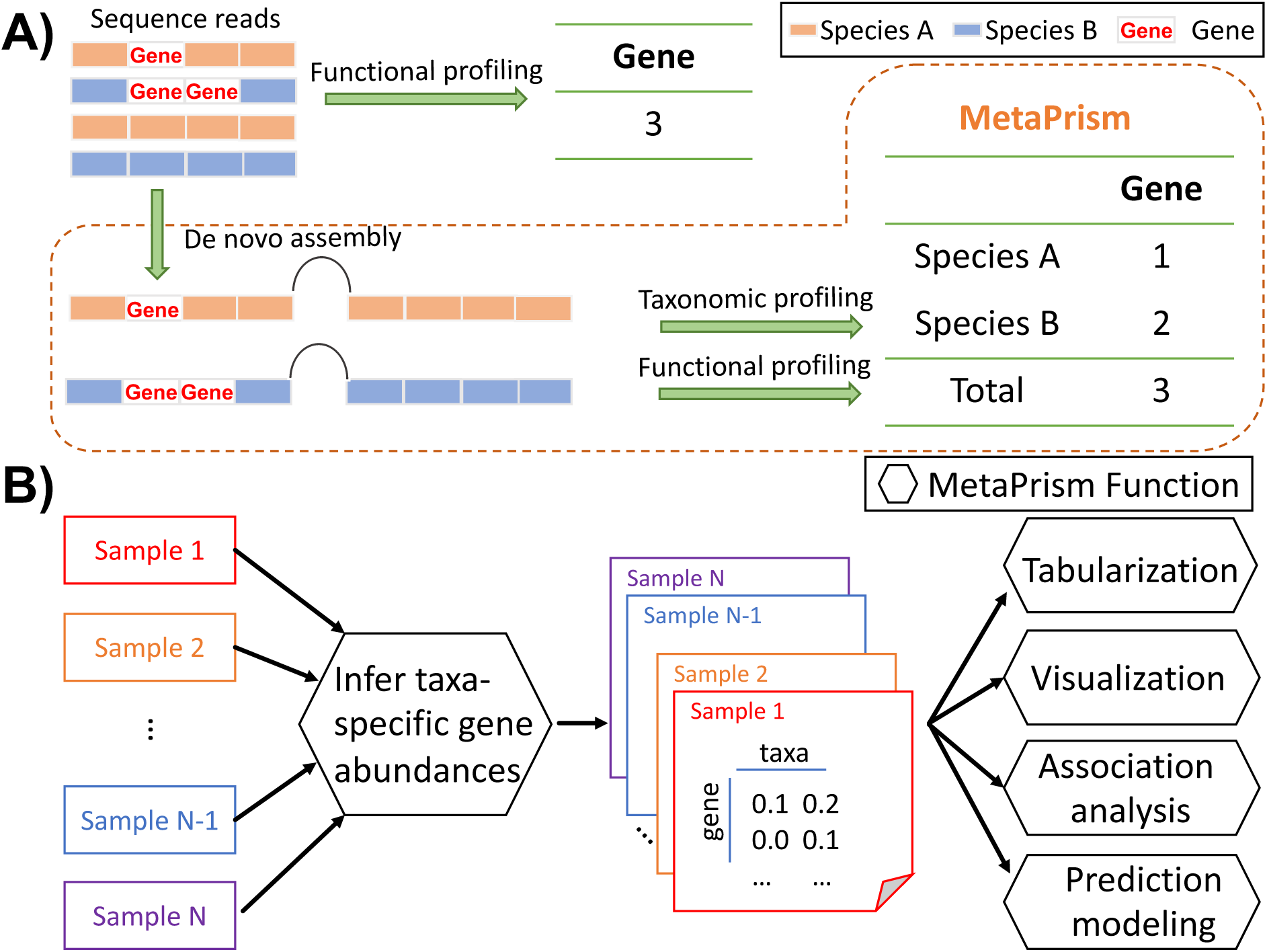
An illustration of the algorithm and the functions in MetaPrism. A) Illustration of the MetaPrism algorithm to infer taxa-specific gene abundances. Function profiling alone infers that three reads are mapped to a gene, but cannot provide further taxonomic information. MetaPrism can estimate two reads are from species A and one read is from species B; B) An overview of the joint analysis workflow in MetaPrism. Hexagons represent functions in MetaPrism.

First, we perform *de novo* assembly for each sample using metaSPAdes [8] with all metagenomic sequence reads to obtain long contigs. As these contigs are much longer than sequence reads, that allows for accurate taxonomical and functional profiling.

Second, we identify the taxonomy of these contigs. All the contigs are aligned to a large reference database of more than 4,000 bacterial genomes using centrifuge [9]. Ambiguous alignments will be filtered out from the subsequent analysis.

Third, we identify genes and their locations from the contigs. We detect the open reading frames from the contigs, translated the nucleotide bases to amino acids, and aligned them using DIAMOND [10] to a protein database. To comprehensively investigate all bacteria genes, either KEGG protein databases that include protein sequences from KEGG orthologue genes [11] or KFU (KEGG orthology with UniProt protein sequences) [4], can be utilized. By default, we required minimum coverage of 0.8 to ensure good protein alignments.

Lastly, we calculate and normalize gene abundance within-sample. We align metagenomic sequence reads to the contigs using BWA [12], and count the number of aligned reads located in the genes of interest. We calculate the read depth normalized by contig length, and this quantity is denoted as mean depth to represent the gene abundances. Larger numbers often indicate higher gene abundance. Other abundance statistics, such as FPKM (Fragment Per Kilobase of transcript per Million reads) or depth per genome (normalized read depth per taxa genome length), are also provided.

Through the above steps, the gene abundances are associated with taxonomy information. To assess the accuracy of these estimations, we conducted a simulation study. First, we selected 115 bacterial species with complete reference genomes and downloaded their sequences from NCBI FTP (ftp://ftp.ncbi.nlm.nih.gov/genomes/genbank/bacteria). Then we simulated shotgun metagenomic sequencing reads and generated at 10X coverage to resemble typical read length from the Illumina (100 bp) using ART [13] “art_illumina --in bacteria_complete_reference_genome.fasta --len 100 --fcov 10 --mflen 200 --sdev 50 --noALN --out bacteria_complete_reference_genome.illumina”. We re-assembled the simulated metagenome sequences using metaSPAdes [8]. To evaluate the gene abundances calculated by different methods, we compared MetaPrism to our previous program named FMAP using translation alignment (BLASTX), as our previous approach was shown to accurately report gene abundances [4]. The true abundances were determined by the KEGG ortholog (KO) abundance of the genes in the reference genomes by aligning them to the KEGG protein database using DIAMOND [10]. In **Figure 2**, The scatterplot visualized the true abundances (x-axis) and the estimated abundances (y-axis), and MetaPrism showed higher correlation (correlation coefficient = 1.000) compared to FMAP (correlation coefficient = 0.985).

**Figure 2.**
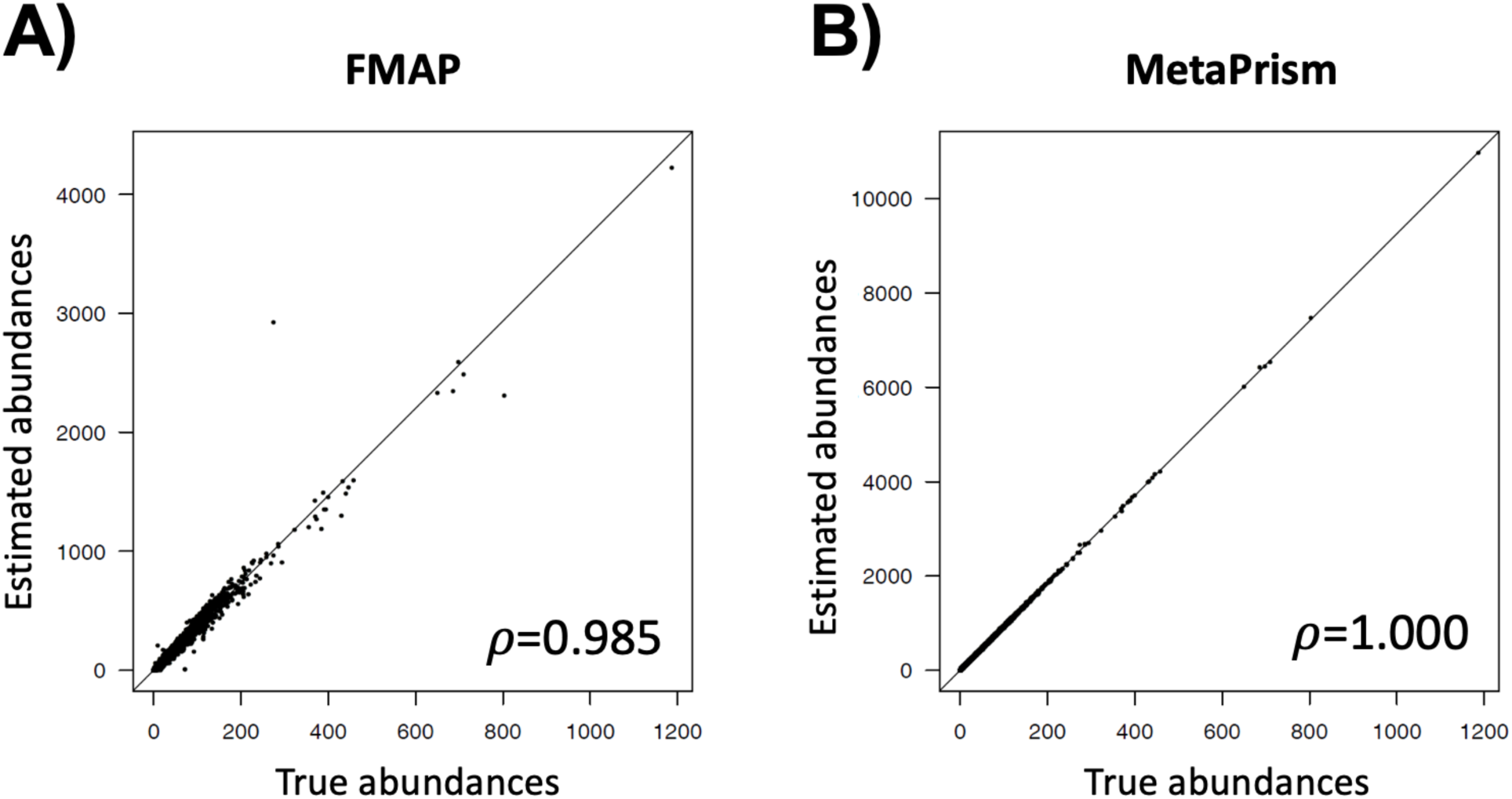
Comparison of gene abundances reported by FMAP and MetaPrism. We used simulations to compare the estimated gene abundances using FMAP and MetaPrism. The Pearson correlation coefficients between true abundances and the software estimated abundances were listed on the bottom right.

In brief, we simulated metagenomic sequence reads from known species and inferred the gene abundances using FMAP (Kim, et al., 2016b) and MetaPrism. This benchmark showed that gene abundances inferred by MetaPrism were accurate and achieved the highest correlation between inferred abundances and true abundances (**Figure 2)**.

Based on these joint features, MetaPrism provided the following downstream joint analysis functions (demonstrated in **Figure 1B**): 1) tabularize the abundances of these features (MetaPrism_table.pl); 2) visualize the features in heatmaps (MetaPrism_heatmap.pl); 3) compare the taxa-specific genes abundances across different experimental conditions such as case-control studies (MetaPrism_compare.pl); 4) indicate which features may serve as potential biomarkers in a prediction model (MetaPrism_predict.pl). A list of available functions, command line, and major customization options in MetaPrism are listed in **Table S2**.

## Results

The gut microbiome plays an important role in modulating immune checkpoint therapy [14]. Here we demonstrated a joint analysis using MetaPrism to build a therapy response prediction model. We collected stool samples of 12 melanoma patients before anti-PD1 (pembrolizumab) therapy and performed metagenomic sequencing. 6 patients responded to the therapy and 6 did not.

Starting from the metagenomic sequence reads, we performed quality control, including removal of human contamination as previously described [14]. Then all remaining sequence reads were processed in MetaPrism (detailed analysis steps provided in **Supplementary Texts**). On average, MetaPrism inferred the taxonomy and gene features for 1.2 billion reads per sample. Next, MetaPrism normalized the reads within samples by reporting the mean depth per assembled contig. The taxa-specific gene abundances were ranked using a random forest model with leave-one-out cross-validation. This prediction model reached 69% accuracy to predict the immunotherapy responses, which is higher than the accuracy using taxa features alone (54%) or gene features alone (62%). Furthermore, it detected four joint features with variable importance greater than 50%. MetaPrism visualized these abundances with red to green colors representing the depth values (**Figure 3**). The most important feature is the K00826 gene (branched-chain amino acid aminotransferase) from the genus Eubacterium (**Table 1**). It is a novel joint feature that may serve as a potential biomarker for cancer therapy.

**Table 1.** Prediction models and performances for taxonomical analysis, functional analysis, and joint analysis. We tabularized the details of prediction models used in three types of analyses and their prediction performances.

|  | Taxonomic | Functional | Joint |
| --- | --- | --- | --- |
|  | profiling | profiling | profiling |
| **Model** | Random forest | Random forest | Random forest |
| **Number of trees** | 500 | 500 | 500 |
| **Number of features** | 1,048 | 5,227 | 62,086 |
| **Top features (if variable importance > 50%) #** |  |  |  |
| **1<sup>st</sup> feature** | Chondromyces (100) | K07705 (100) | K00826 Eubacterium (100) |
| **2<sup>nd</sup> feature** | Roseateles (65) | - | K03205 Streptococcus (89) |
| **3<sup>rd</sup> feature** | - | - | K01006 Flavonifractor (81) |
| **4<sup>th</sup> feature** | - | - | K06187 Thermoanaerobacter (74) |
| **Accuracy\*** | 53.8% | 61.5% | 69.2% |

#: The variable importance values are listed in parentheses
\*: Prediction accuracy was evaluated using leave-one-out cross-validations

**Figure 3.**
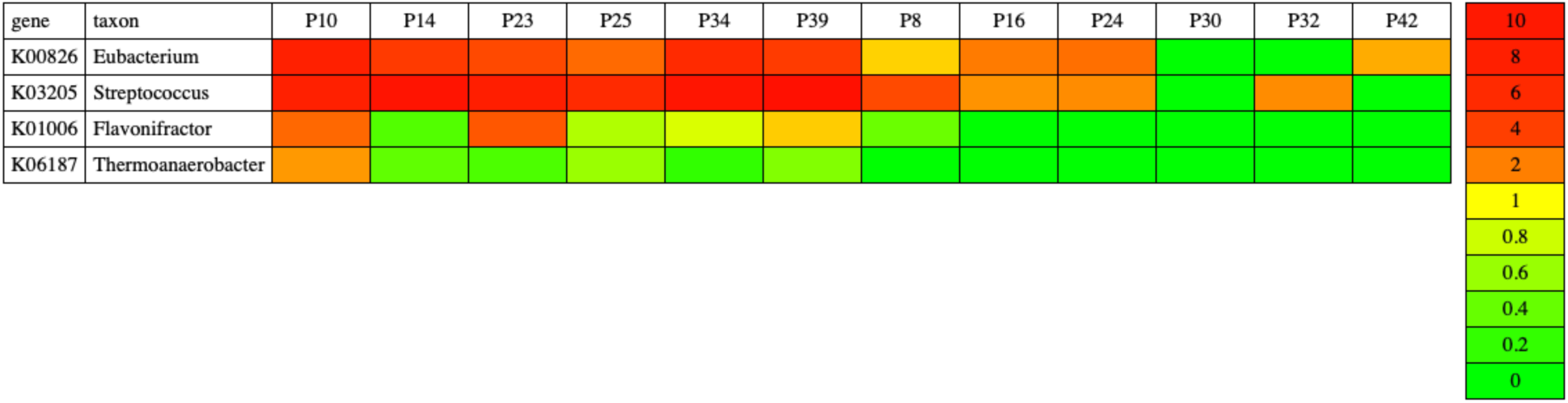
Heatmap of joint features for predicting immune checkpoint therapy response. We used MetaPrism_heatmap.pl to visualize four joint features (taxa-specific gene abundances) in the immune checkpoint therapy study. The colors from red to green represent the increased gene abundances, the mean depth normalized by the contig lengths. P10, P14, P23, P25, P34, P39 are patients who respond to the therapy; P8, P16, P24, P30, P32, P42 are patients having progressive outcomes.

In terms of computation, all the above analyses can be accomplished on a standard computation cluster (e.g., 128GB memory with 2 GB hard drive space per sample).

## Discussion

We present a novel bioinformatics tool, MetaPrism. It implements functions to quantify the joint features (both taxonomic and functional) from metagenomic sequence reads, as well as other functions for downstream data analyses. We demonstrate that the joint features can provide novel insights to understand the microbial role in a cancer immunotherapy study.

MetaPrism is flexible and can be customized. For example, to study species-specific antibiotic resistance genes (ARGs), a reference protein database with ARGs, such as ARDB [15] or CARD [16], can be used. MetaPrism can infer taxa-specific ARGs, thus enabling joint resistome profiling. In a GVHD study, with the interests to study the patients’ resistome, we performed the analysis using the ARDB in MetaPrism and found increased abundances of antibiotic-resistance genes (e.g., *mdtG, AcrA, AcrB*, and *TolC*) in *Klebsiella* and *E. coli* in the GVHD patients compared with the abundances in non-GVHD patients. This finding may hint optimal antibiotic prescription for better management of GVHD.

MetaPrism characterizes the joint features based on the contigs that are *de novo* assembled from metagenomic sequence reads. This is a distinct feature compared with other software. For example, HUMAnN2 used a tiered search strategy that relied on a curated reference database for organism-specific genes[3]. As human microbiome contains trillions of microbial genes, reference databases can be inadequate to enumerate the organism-specific genes. Thus we designed the MetaPrism to reduce the dependency on curated reference databases. The tradeoff for this decision is that MetaPrism requires more computational resources for the *de novo* assembling step.

In all, MetaPrism is free and useful software to facilitate joint analyses and it is suitable for general microbiome studies. Researchers can expect MetaPrism to quantify species-specific gene abundances and use these interpretable features in association studies and prediction tasks.

## Availability and requirements

**Project name:** MetaPrism

**Project home page:** https://github.com/jiwoongbio/MetaPrism

**Operating system(s):** Platform independent

**Programming language:** Perl, R

**Other requirements:** None

**License:** GNU GPL

**Any restrictions to use by non-academics:** None

## Declarations

### Availability of data and materials

The metagenomic shotgun sequence dataset are available from the NCBI BioProject PRJNA397906. The treatment responses for the 12 patients as well as the analysis codes were available in the **Supplementary Texts**. The source codes of MetaPrism software is available at: https://github.com/jiwoongbio/MetaPrism. That resource contains the software requirements, usage example and documentations for all MetaPrism components (e.g., download bacterial database, quantify species-specific gene abundances, build association models and prediction models, tabularize results and visualize results in heatmap).

### Authors’ contributions

JK, SJ, and XZ conceived of the project and wrote the first draft of the manuscript. GX, YX, AY, and XZ coordinated and oversaw the study. QL, DL provided critical inputs for the study. JK and SJ developed the software and associated databases. All authors contributed to the review of the manuscript before submission for publication. All authors read and approved the final manuscript.

### Ethics approval and consent to participate

Not applicable.

### Consent for publication

Not applicable.

## Acknowledgement

We thank Jessie Norris for her comments on the manuscript.

